# AAV Vector Transduction Restriction and Attenuated Toxicity in hESCs via a Rationally Designed Inverted Terminal Repeat

**DOI:** 10.1101/2022.06.30.498264

**Authors:** Liujiang Song, Nolan J. Brown, Jacquelyn J. Bower, Richard J. Samulski, Matthew L. Hirsch

## Abstract

Recombinant adeno-associated virus (rAAV) inverted terminal repeats (ITRs) induce p53-dependent apoptosis in human embryonic stem cells (hESCs). To interrogate this phenomenon, a rationally designed ITR (SynITR), deleted for p53 binding sites was evaluated for vector production and gene delivery. While SynITR genomes were decreased for transgenic genome replication compared to wtITRs, similar production titers indicated that replication is not rate-limiting. Packaged in the AAV2 capsid, wtITR and SynITR vectors demonstrated similar transduction efficiencies of human cell lines with no differences in reporter kinetics. Following rAAV2-wtITR infection of hESCs, rapid apoptosis was observed, in contrast, rAAV2-SynITR infection resulted in attenuated hESC toxicity with cells retaining their differentiation potential. While hESC particle entry and double stranded circular episomes was similar for the ITR contexts, reporter expression was significantly inhibited from transduced SynITR genomes. Infection of hESCs induced γH2AX in an ITR-independent manner, however, canonical activation of p53α was uncoupled using rAAV-SynITR. Further hESC investigations revealed 2 additional novel findings: i) p53β is uniquely and constitutively active, and ii) rAAV infection, independent of the ITR sequence, induces activation of p53ψ. The data herein reveal an ITR-dependent rAAV transduction restriction specific to hESCs and manipulation of the DNA damage response via ITR engineering.

## Introduction

Although adeno-associated virus (AAV) is widely considered a non-pathogenic and non-autonomous virus, data also report the opposite; namely that AAV transduction can induce cell death and/or that AAV is competent for autonomous replication (1-7). Interestingly, these seemingly uncharacteristic behaviors of AAV appear to manifest in particular, perhaps natural, contexts including in superinfected cells and in less differentiated cellular states such as cancers and stem cells (8-10). In several of these instances, the ITR sequence itself has demonstrated the capacity for cellular toxicity including in models of adenocarcinoma and early pregnancy (1, 2, 11). This AAV ITR sequence has now been administered by various routes to thousands of human patients as an inherent component of the drug formulation of all recombinant AAV gene therapies.

The AAV ITRs are largely conserved in sequence and function across most AAV serotypes (12, 13). The AAV serotype 2 ITRs (referred to as wtITRs herein) were the first ITRs cloned into a plasmid and continue to be the most employed for the overwhelming majority of applications for both experimental and clinical gene therapy (14). Albeit relatively small, the 145nt wtITR sequence serves many functions including: i) the origin of replication, ii) the capsid packaging signal, iii) telomere, iv) priming of second strand synthesis, v) episome circularization and persistence, vi) transcriptional activity and enhancer functions, and vii) site-specific integration on human Ch19 (15). Several of these functions, rely on the fact that these telomere-like sequences have repeating nucleotide patterns, and as terminal inverted repeats, are predicted to fold into complex structures occluding free ssDNA ends from host exonuclease activity while providing duplexed binding sites for endogenous and host factors. Additionally, the wtITR sequence is inherently GC rich, which is a feature of human replication origins and perhaps consequently transcriptionally active regions (12, 16). The repeating pattern of precisely spaced G nucleotides on ssDNA exhibited by the wtITR is predicted to adopt a G-quadruplex formation, which are common amongst viruses and are also features of replication origins and human telomeres (17, 18). Collectively, these attributes of wtITRs induce a cellular DNA damage response (DDR) upon transduction, the consequences of which appear to be cell type specific.

Human embryonic stem cells (hESCs), derived from the inner cell mass of the blastocyst, are totipotent, meaning they can differentiate into any cell type of the human body and presumably can still generate a viable human in theory. Perhaps because of their ability to influence all progeny cells, it is widely appreciated that hESC’s tolerance to insults, especially genotoxic stress, is very low. In response to DNA damage, hESCs display altered cell cycle responses that culminate in rapid apoptosis (19). A similar mechanism is reported when AAV vectors are the “genotoxic” stress, and that following hESC transduction, p53-dependent apoptosis occurs in early G1 phase of the cell cycle (2, 3). No observed cellular response to empty AAV capsids, but apoptosis upon ITR DNA microinjections implicated the causative agent as the nucleotide sequences within the ITR (2). Interestingly, a discrete G-rich ITR region induced hESC apoptosis upon microinjection while the C-rich complementary strand (also wtITR sequence) did not; these data along with oligonucleotide associations observed by native gel electrophoresis, suggest the potential for a G-Quadruplex induced DNA damage response (2).

Given these and other reports of AAV ITR DDR in stem cells, it was hypothesized that alteration of the ITR sequence would alter the toxicity and potentially other aspects of AAV vectorology (4, 9). Therefore, a synthetic ITR (SynITR) in which putative p53 binding sites were ablated by nucleotide substitution was investigated. Data herein demonstrate its competency for vector production despite a decreased replicative capacity compared to the wtITR. Additionally, SynITR vectors mediated transduction of several human cell lines at similar efficiencies and kinetics as wITR-based vectors. Interestingly, AAV2-SynITR vectors attenuated toxicity following hESC transduction compared to wtITR vectors. This was despite similar intracellular vector genome accumulation and double strand (ds) circular episome formation among the vector contexts. However, transgenic reporter transcription and subsequent reporter activity was dramatically decreased for AAV-SynITR vectors, suggesting an ITR-based host factor restriction of rAAV in hESCs. At the mechanistic level, the SynITRs induced an altered DDR compared to the wtITR when packaged in the same capsid. The work herein highlights the ability to engineer ITR sequences competent for all aspects of production with the capacity to alter the cellular DDR upon transduction, perhaps resulting in a potentially safer methodology for hESC transduction and perhaps stem cells in general. Additionally, novel insights regarding unchallenged hESC physiology and rAAV vector induction of p53ψ were noted.

## Results

### Design/Engineering of SynITR

In previous work we and others reported that AAV vector transduction of hESCs induced p53-dependent apoptosis (2, 3). Specifically, the AAV ITR serotype 2 (wtITR) sequence itself was sufficient for apoptosis highlighting a lethal DNA damage response (DDR) in undifferentiated cells to a viral telomere employed in human gene therapy. Upon sequence examination of the 145 nucleotide wtITR sequence, many predicted transcription factor binding motifs were identified, including 6 putative p53 binding sites (Fig. 1A). This led to the hypothesis that ablation of these putative binding sites would alter the DDR in hESCs and therefore allow AAV vector transduction without toxicity. Therefore, the guanine-cytosine rich p53 putative binding motifs in the B/C regions of the wtITR were substituted with adenine-thymine nucleotides while retaining the native A/D region (termed Synthetic or SynITR, Fig. 1). Compensatory mutations were engineered to maintain predicted stem folding (Fig. 1B). Sequence analysis of SynITR reveals reduced GC content, which consequently decreased the free energy of its overall DNA folding, as well as a decreased potential for G-quadruplex formation compared to the wtITR (Fig. 1b). Furthermore, the sequence modifications ablated many other predicted host factor binding sites in addition to those of p53 (Fig. 1 and website). Initially constructed by triple ligation in the reported “DD” CMV-GFP plasmid context (20), and propagated in recombination deficient SURE cells, the integrity of the SynITR was stable as determined by frequent DNA sequencing and by enzymatic digestion after each plasmid preparation using an unique engineered BsaBI site (20).

**Figure 1.**
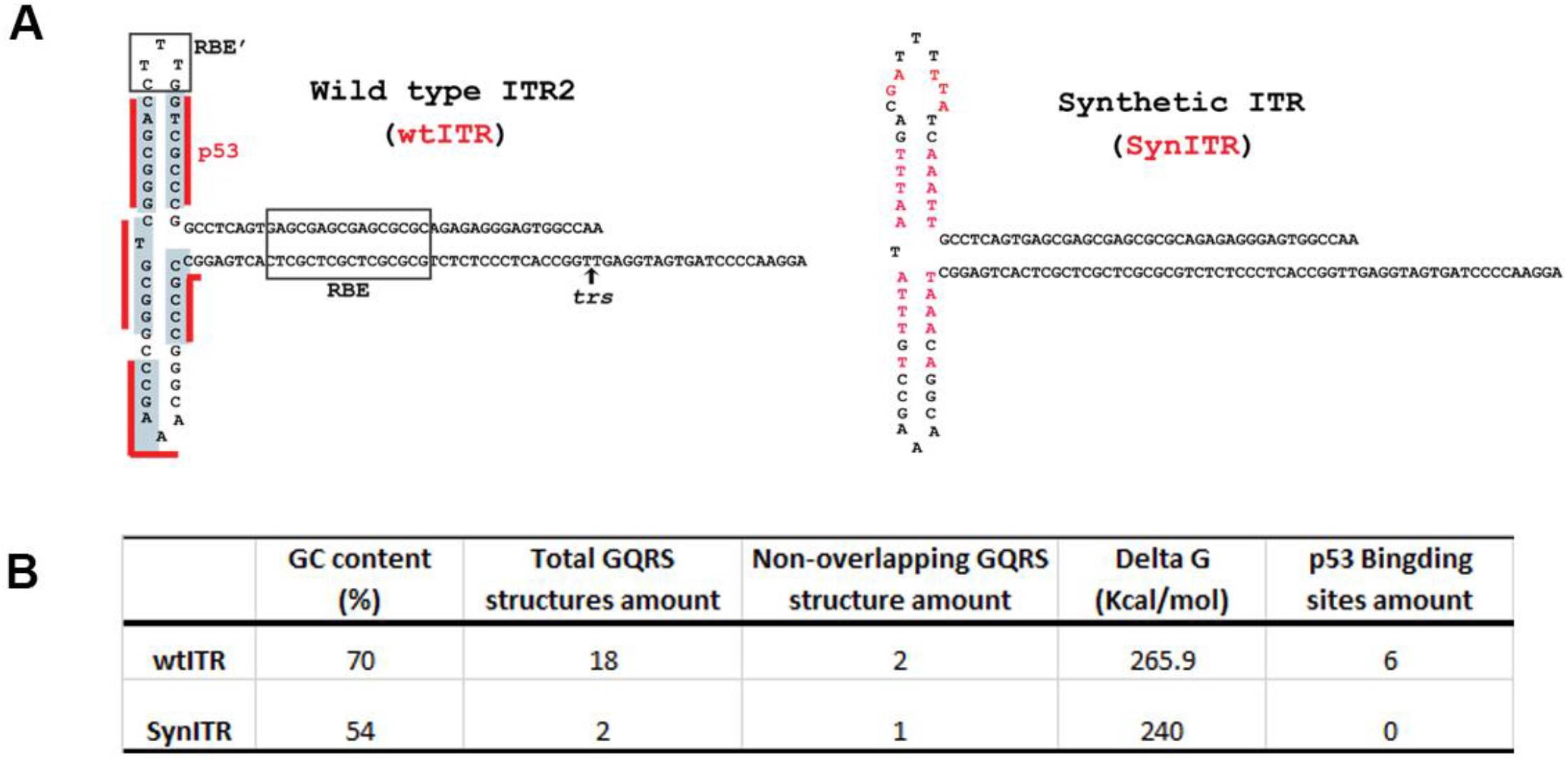
A. wtITR and SynITR sequence with putative p53 binding sites overlined in red and nucleotides unique to SynITR in red font. B. Sequence analysis. Web-based tools were utilized for GC content, prediction of GQRS, free energy, and the potential p53 transcription factor binding sites.

### SynITR plasmid is decreased for replication compared to wtITR

First, the ability of the SynITR to act as an AAV replication origin in concert with the AAV Rep serotype 2 (Rep2) protein was investigated. HEK293s were transfected with pXR2, pXX680, and the wtITR or SynITR plasmid. Hirt DNA was extracted at 48 hours post transfection, subjected to DpnI digestion, and then resolved on a neutral agarose gel followed by Southern blot analysis using a radiolabeled GFP probe (Fig. 2). Following DpnI digestion to remove the bacterial amplified DNA, samples from wtITR transfection (Fig. 2, lane 4) revealed a nascent banding profile that is characteristic of canonical replication (20). In contrast, only a weak replicative monomer species was observed demonstrating that the SynITR is decreased for plasmid resolution and/or Rep2 mediated replication (Fig. 2, lane 5). When the wtITR and the SynITR plasmids were co-transfected in a competitive assay, replication of only the wtITR substrate was observed (Fig. 2, lane 6).

**Figure 2.**
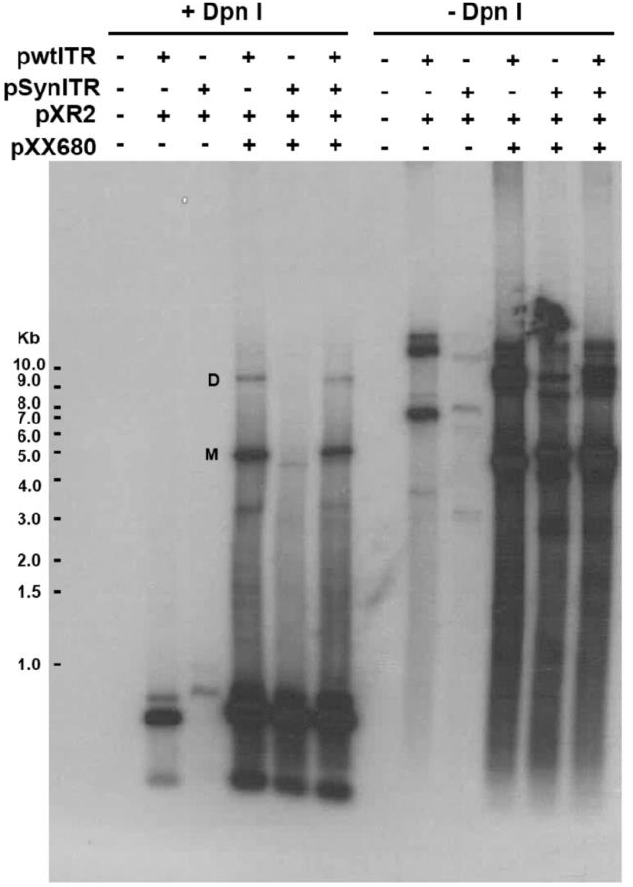
Examination of vector-genome replication in a three plasmid co-transfection reaction. HEK293 cells were co-transfected with the pXR2, pXX680, and the ITR-containing AAV plasmid. Hirt DNA was harvested at 48 h post transfection, digested with or without Dpn I and analyzed by Southern blot using p32 labeled probe. Duplex monomer (M) and duplex dimer (D) replication form were indicated in the image.

### SynITR plasmids produce rAAV vectors

As it is reported that AAV vector genome replication, in itself, is not a rate limiting step in vector production (21), the ability of the SynITR replication substrate to be packaged in the AAV capsid was exhaustively investigated in parallel with an wtITR plasmid (both contain a CMV-GFP cassette) using a standard triple transfection production protocol (22). First, the genome-containing particle titer was determined by probe-based qPCR as described using multiple probe/primer sets as well as by Southern dot blot analysis. Consistently across assays, the SynITR yielded approximately a 3-fold higher titer of DNAse I resistant viral genomes than the wtITR2 preparation (Fig. 3A, B, S2A). Western blot analysis, loaded by the qPCR titer and using the B1 antibody, showed no obvious difference regarding the capsid protein ratio of VP1:VP2:VP3 between the wtITR and SynITR preparations, however less overall abundance of these proteins was observed for in the SynITR group (Fig. 3C). Therefore, AAV2 capsid to viral genome ratio was analyzed via Western dot blot with the A20 antibody which recognizes the intact AAV2 capsid. While the same titer of vector genomes was loaded, there was a consistent stronger signal for the wtITR preparations suggestive of more empty capsids in that preparation (Fig. 3D). At the genome level, alkaline gel electrophoresis confirmed the integrity of the packaged SynITR genome and sequencing confirmed that the SynITR was packaged within the capsid (Fig. 3E and S2B; a few instances of sequence ambiguity were also noted and are commonly reported for the wtITR (Fig. 3E). Additionally, the inadvertent packaging of the Rep sequence genome during vector production was analyzed in the preparations by qPCR with no significant difference observed between the wtITR or Syn ITRs preparations (Fig. S2C)(23).

**Figure 3.**
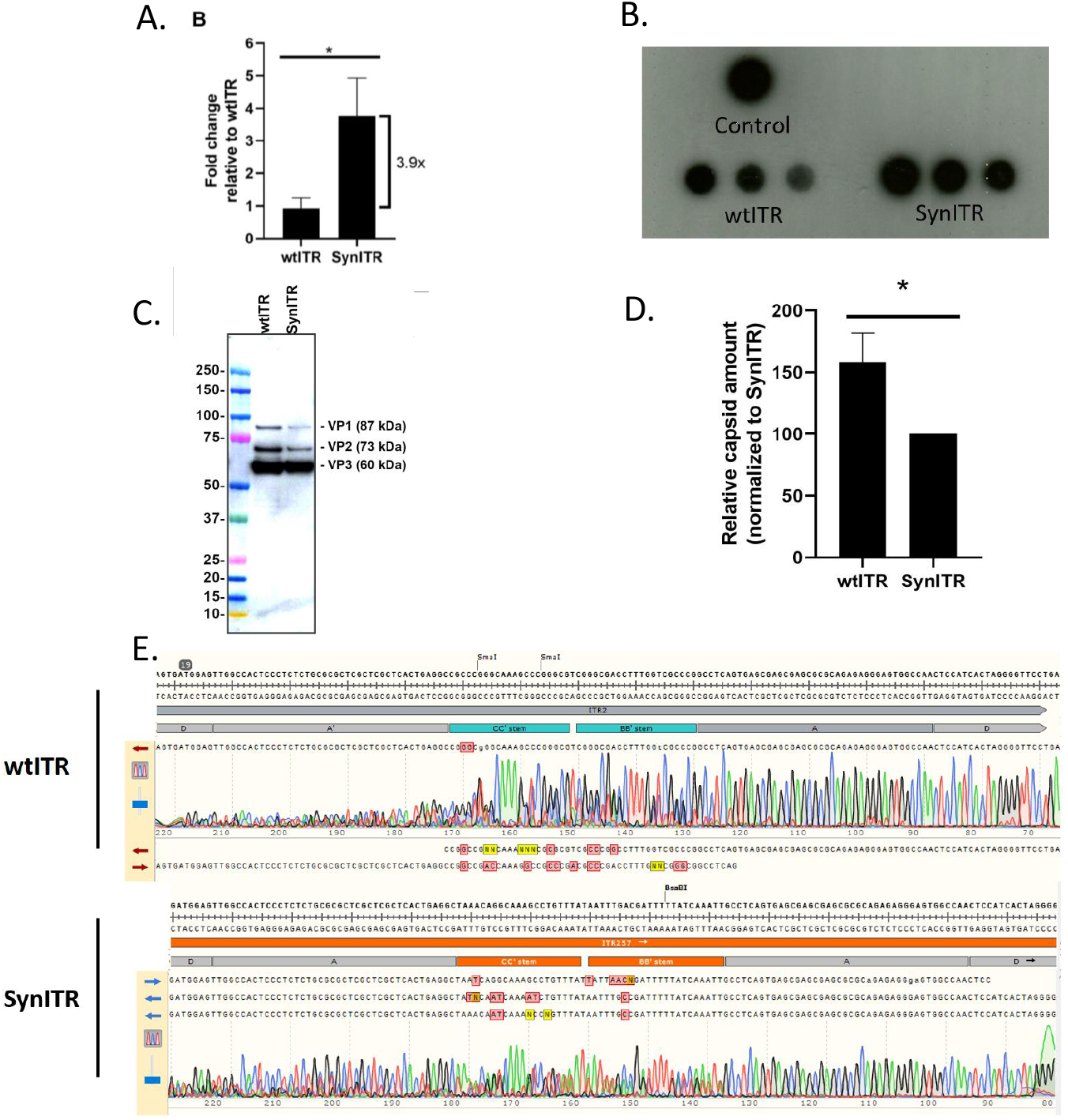
Comparison of AAV production and vector characterization. A, Quantification of the vector genome yield by probe-based qPCR. *p<0.05. B. Virus titering by Dot blotting. C, Representative AAV capsid western blot using the B1 antibody. D, Representative AAV capsid dot blot using A20 antibody that only recognizes the intact AAV capsid. E, Densitometry analysis of capsid yield by Image J. F. Viral genome sequencing.

### AAV-SynITR mediates transduction of differentiated human cell lines

AAV2 vectors harboring the SynITR (AAV2-SynITR) were evaluated for transduction in HEK293s, human liver Huh7 cells, and h-TERT immortalized diploid human fibroblasts (NHFs). Using 10,000 vg/cell, GFP+ cells were visualized by microscopy and tallied by flow cytometry 72_hr post-vector addition. The results demonstrated that AAV2-SynITR vectors were competent for transduction at levels similar to wtITR vectors in the tested contexts (Fig. 4A, B). Additionally, cell viability following vector transduction with either wt or Syn ITR flanked genomes demonstrated no AAV vector induced toxicity in these tested cell lines (Fig. 4C).

**Figure 4.**
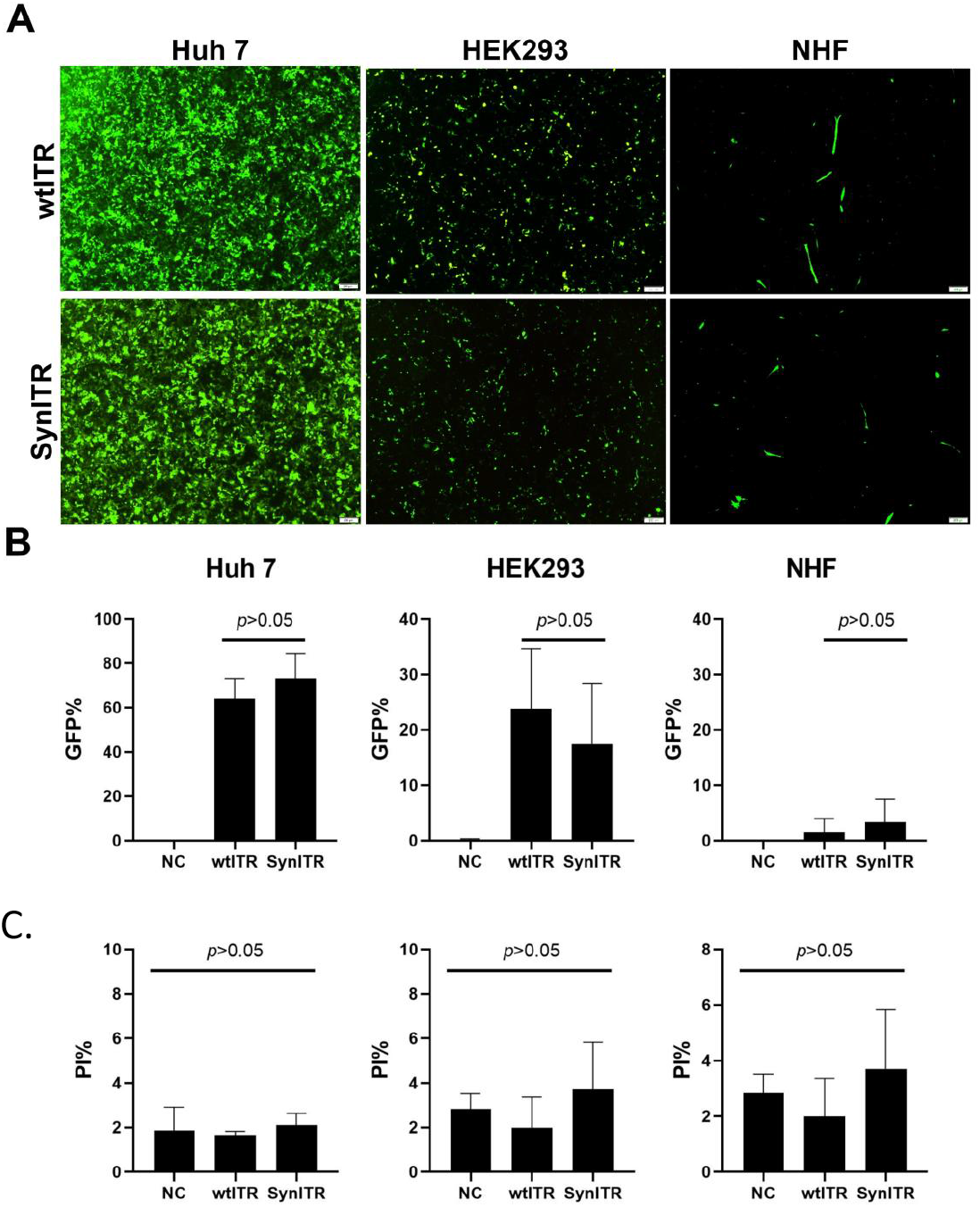
SynITR does not significantly alter transduction efficiency of a CMV-GFP reporter in differentiated lines. Huh 7, HEK293 and NHF cells were transduced with 10,000 vg/cell. Transduction efficiencies were examined at day 3 by microscopy (A); flow cytometry for %GFP+ cells, (B) and %PI+ for cell viability (C).

### AAV2-SynITRs attenuate toxicity and are restricted for transduction of hESCs

Next, the central hypothesis of this work was tested by infecting WiCellH9 hESCs with AAV2 capsids packaged with either wtITR or SynITR flanked single strand transgenic cassettes. As previously reported, ssAAV ITR2 vector transduction results in rapid hESC apoptosis confirmed by microscopy and Western analysis of cleaved PARP-1 (Fig. 5A-C)(2, 3). In contrast, SynITR vectors induced only minimal toxicity when exposed to hESCs (Fig. 5A-C). Despite the equivalent number of intracellular genomes as wtITR based vectors (Fig. 5E), and similar transduction efficiencies in other cell types (Fig. 4), GFP+ cells and gfp cDNA were significantly decreased for the SynITR-treated hESCs (Fig. 5D, E). A previously reported AAV “uncoating assay”, which measures the abundance of transgenic circular dsDNA, was executed following hESC exposure to either of the ITR vector contexts. The results indicate that approximately 0.4% of vector genomes were T5 exonuclease resistant with no significant differences between wtITR and SynITR treated cells (Fig. S2A). Interferon β, an innate immune effector often activated following AAV infection, and having the capacity for apoptotic induction was also investigated (24). 24 hours following rAAV exposure, RT-qPCR analysis demonstrated no significant difference between untreated hESCs and groups receiving rAAV with wtITR or SynITRs (Fig. S2B).

**Figure 5.**
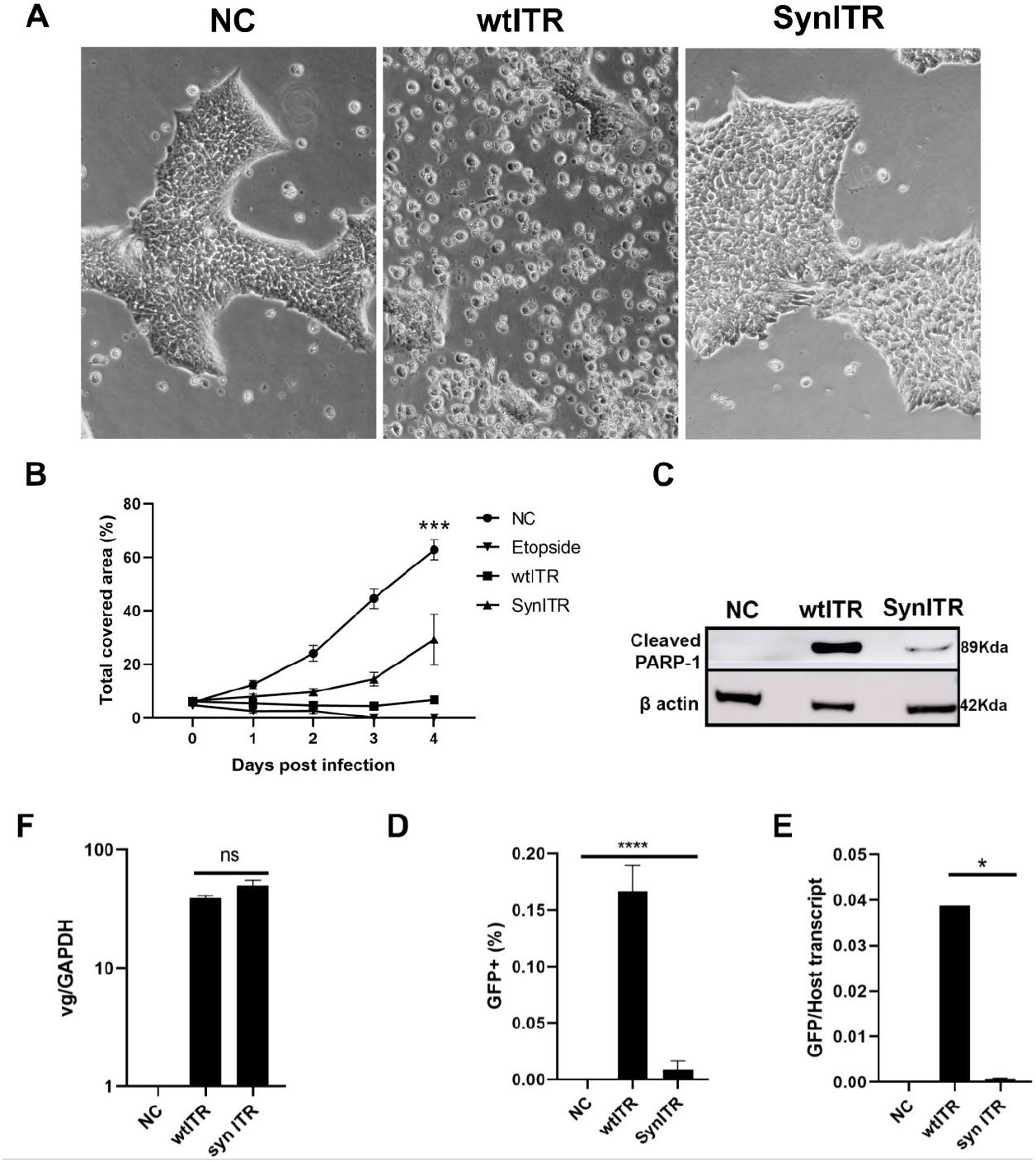
AAV ITR sequence affects the transduction efficiency and toxicity in hESC WiCell H9 cells. H9 cells were propagated under feeder-independent conditions, seeded in Matrigel treated 24-well plates, and cultured with StemFlex medium to allow colony formation. A, Representative images of H9 cells transduced with AAV. Severe cell death was observed approximately 24 hours post infection by wtITR vector. B, Colony growth examined by measuring the plate surface area covered by clones. C, Western blot detection of apoptosis. **D**. Transduction efficiency by wtITR and SynITR vectors in human embryonic stem cell H9 cells examined by flow and RT-QPCR (E). AAV with wtITR resulted in significant higher transgene expression in H9 cells. F. No significant differences were observed in the number of viral genomes/cell.

Next, it was determined if hESC differentiation following exposure to wtITR2 or SyntITR vectors would alter the observed wtITR2 toxicity (Fig. 6). In these experiments, hESCs were exposed to either AAV2 vector ITR context and then embryoid body (EB) formation assay was initiated. Five days later, significantly fewer EBs were derived from hESCs exposed to wtITR vectors (Fig. 6). In contrast, infection of hESCs with SynITR vectors did not alter the potential for EB formation compared to the negative control (Fig. 6).

**Figure 6.**
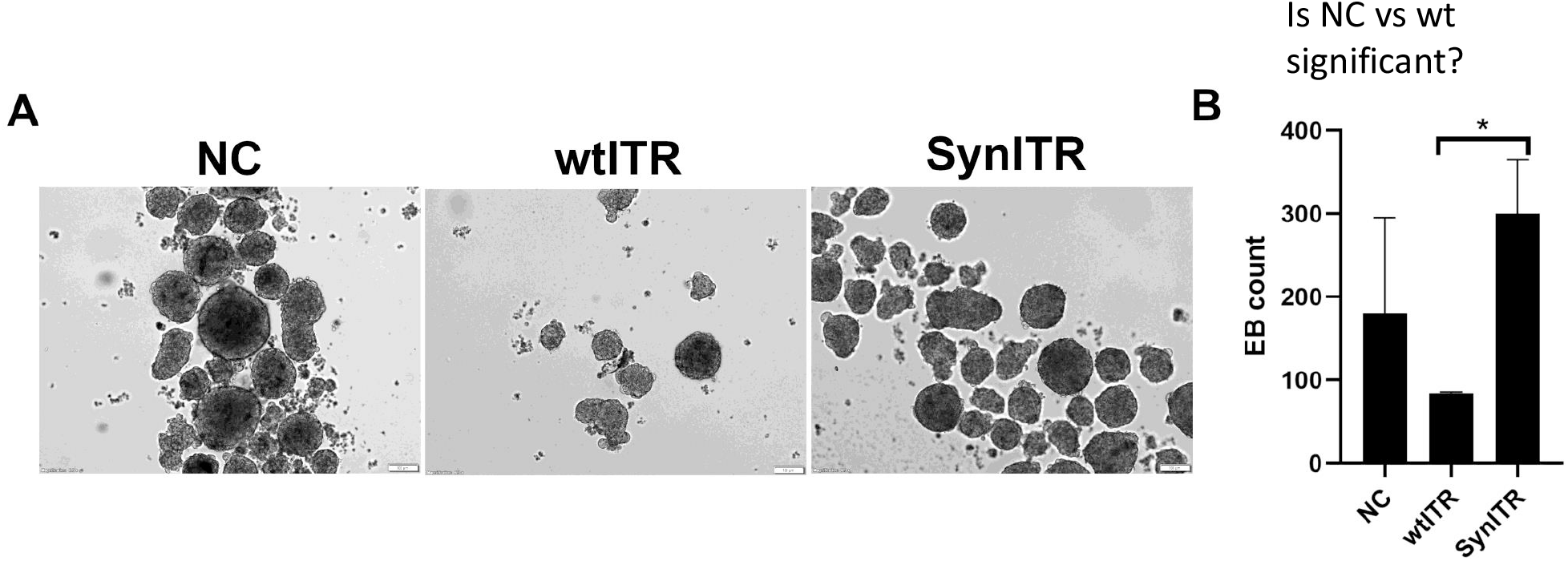
AAV transduction affects embryoid body formation. The EB formation assay was performed using the DMEM-F12 containing 20% KnockOut™ Serum Replacement with the addition of a ROCK-inhibitor and other nutrients as stated in the methods. EB plates were scanned, and the number of EB spheres were counted on Day 7 culture.

### AAV SynITRs alter the DNA Damage Response in hESCs

In previous reports in hESCs, single-strand and self-complementary AAV vectors activated p53, which was required for vector-induced apoptosis (2, 3). Therefore, total p53 and a marker of p53 activation, phosphorylation of serine 15, was investigated in hESCs transduced with either wtITR or SynITR vectors by Western immunoblot analysis. For these experiments, diploid human fibroblasts, which are well characterized for DDR signaling served as a control cell line. Additionally, hydroxyurea (induction of replication stress) and etoposide (double-stranded break signaling) were used as controls. First, total p53 (using a N-terminal antibody) was increased in hESCs in response to etoposide or infection by wtITR vectors harboring a ss or sc transgenic genome (Fig. 7A). In contrast, SynITR vectors did not increase total p53 abundance following infection by this assay (Fig. 7, DMSO vs. SynITR). This result is consistent with a 30-fold decrease in p53 transactivation of the p21 transcript using SynITR vectors compared to those harboring wtITRs (Fig. 7B). Interestingly, a smaller variant of p53, reported as p53ψ (27kDa) was observed in hESCs treated with rAAV, independent of the ITR sequence, but not in the hydroxyurea or etoposide treated cells (Figs. 7, S4)(25). Additional isoforms of p53 corresponding to β and/or γ were also noted, in both NHFs and the hESCs (Fig. 7). Next, p53ser15 phosphorylation, a commonly used marker of p53 activation, was investigated in response to the different stimuli. Unexpectedly, it was noted that, specifically, in unchallenged hESCs p53γ and/or p53β isoforms (48kDa or 47kDa respectively) were constitutively activated, and no variation was noted upon drug or AAV vector exposure (Fig. 7). RT-qPCR analysis with p53γ or p53β specific primers noted remarkably increased p53β cDNA compared to γ, suggesting the observed band is p53β-Ser15-P. The high levels of activated p53β appeared specific to hESCs when compared to NHFs (Figs. 7A and S4). Although, full length p53α demonstrated robust p53-ser15 phosphorylation in hESCs upon etoposide treatment, p53α activation (as inferred by this specific marker) was noted in the etoposide group and observable in scAAV treated cells at longer exposures (Figs. 7 and S4). However, variations in higher order complexes, reported as p53 homodimer complexes (reviewed in (26)), demonstrated increased abundance in hESCs treated with SynITR, but not wtITR2, vectors (Fig. 7A “II”). Elevation of γ-H2AX abundance was noted in hESCs treated with single strand or self-complementary vectors containing wtITR2 compared to SynITR vectors (Fig. 7).

**Figure 7.**
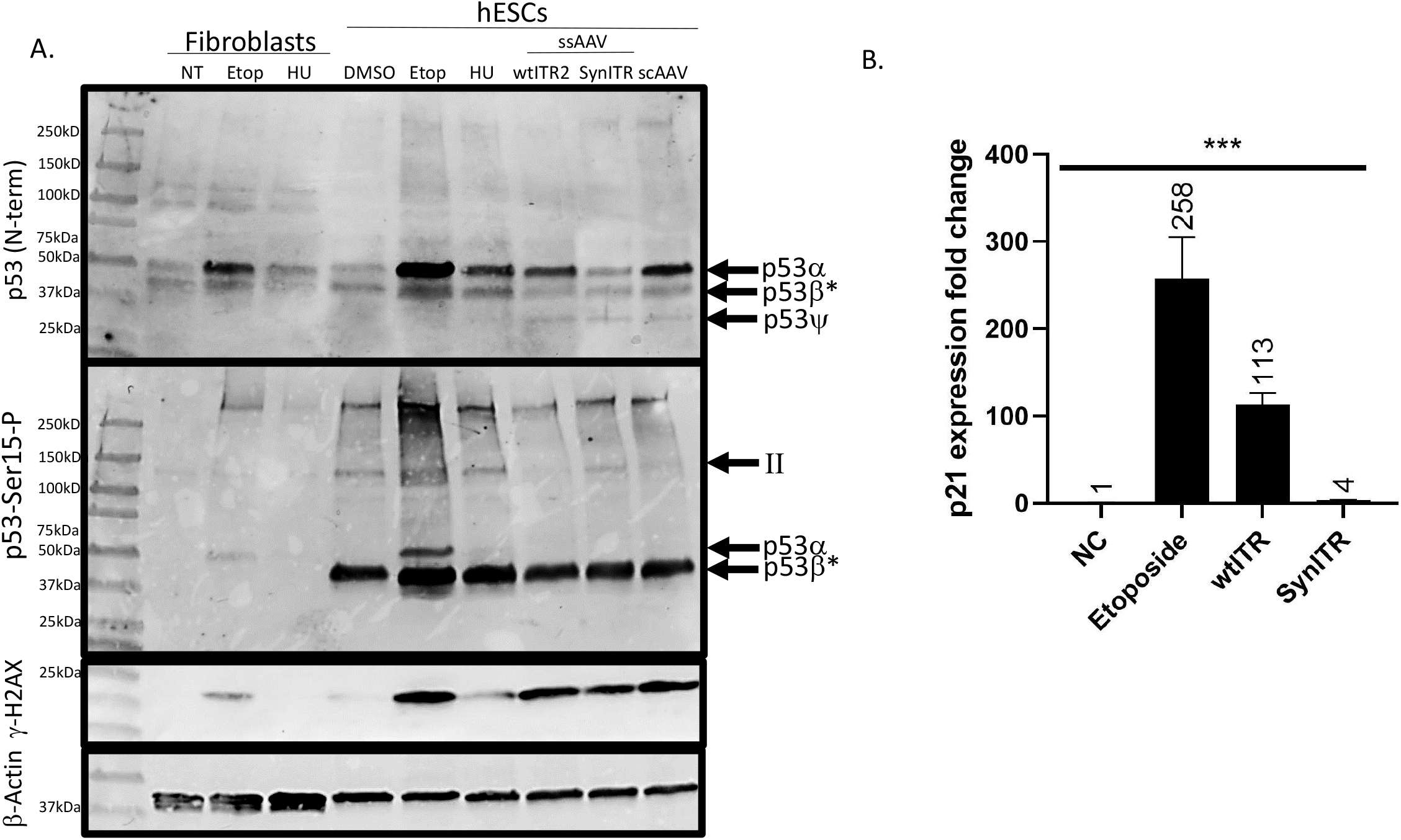
Differential DNA damage signaling by wtITR and SynITR vectors. (A) These experiments using WiCell H9 cells and diploid human fibroblasts treated with etoposide (etop), hydroxyurea (HU), drug vehicle (DMSO), self-complementary (sc) AAV2 with wt ITRs, and single strand (ss) AAV2 with the indicated ITRs. Cellular lysates were analyzed by Western blotting using p53 (N-terminus), p53-ser-15-P (p53 activation), γ-H2AX, or β-actin antibodies. Differential p53 isoforms are labeled. II refers to a predicted p53α homodimer as described in the text. (B) RT-qPCR analysis of p21 expression in H9 hESCs following the indicated treatments. NC = negative control.

## Discussion

The work herein validated the hypothesis that a modification of the ITR altered the DDR in hESCs to decrease vector induced toxicity upon transduction. First, a SynITR 18% dissimilar from wtITR functioned as a substrate for replication in the presence of the wtAAV proteins and adenovirus helper functions (Figs. 1, 2). Albeit decreased for replication, the SynITR resulted in normal to enhanced AAV titer production compared to the wtITR and its packaging was confirmed from the capsid by sequencing (Fig. 3). AAV-SynITR vectors resulted in equivalent transduction efficiencies and kinetics compared to AAV-wtITR vectors in the immortalized or transformed cell lines used herein (Fig. 4). However, an ITR-dependent restriction of transduction was identified which coincided with attenuation of rAAV mediated hESC apoptosis. Based on the collective data from an “uncoating” assay along with the induction of DDR signaling by both wtITR and SynITR vector contexts, it is hypothesized that this ITR-based host restriction is elicited following partial or complete capsid uncoating (Figs. 7, S2; (27). Interestingly, the mechanistic analyses revealed novel insights into the unchallenged state of hESCs, which demonstrate constitutive activation of presumably p53β, in contrast to p53α induction by genotoxic stress (Figs. 7, S3, S4). Another unexpected finding was the induction of p53ψ, a pro-metastatic protein which has only been reported in two murine reports to date, in a manner that was ITR-independent, yet AAV vector specific (Figs. 7, S4; (25, 28). Collectively, the data presented herein provide proof-of-concept for additional rational ITR design to modulate the vector-induced DDR following transduction towards safer and or more efficient rAAV gene delivery to traditionally refractory cell types. Furthermore, preliminary evidence suggests the varied DDR response by SynITRs influences the ability to serve as substrates for gene editing.

AAV vector genome replication from plasmid templates initially requires release (or resolution) from its double stranded circular nature. Then, Rep multimer binding occurs with minimally 2 contact regions within the ITR termed the Rep binding element (RBE), a tetrad-like repeat in the A region serves as the main contact, and RBE’, a sequence found within a predicted small loop of the B/C sequence is reported to orient Rep’s endonucleolytic activity towards the trs (Fig. 1)(29, 30). The binding event along with Rep-mediated helicase activity is thought to result in protrusion of a trs stem then Rep creates a single strand break followed by strand synthesis. Regarding the decreased replication from the SynITR, any one or perhaps a combination of these steps could be less favorable (Fig. 2). However, the RBE sequence itself was maintain along with the native D element (including the trs), suggesting that binding may not be the primary reason for decreased SynITR replication. Interestingly, if it is related to Rep’s binding and nicking activity, then one might speculate the SynITR may also be decreased for specific integration at (or around) the AAVS1 sequence on chromosome 19 (31). Further testing is necessary to determine the precise mechanism.

Given the decreased replication of SynITR genomes, it is of note that it was still able to function for AAV capsid packaging and resulted in titers comparable to those elicited from the wtITR. This observation suggests that under the “pristine” laboratory conditions that AAV vector genome replication is not a rate limiting step during controlled production and is consistent with conclusions of a previous report (21). Another production aspect tested herein was the inadvertent packaging of wtAAV sequences that is reported to contaminate the final preparation across production modalities (23). Despite alterations in the B/C elements of the AAV ITR, wtAAV genome contamination was similar for both wtITR and SynITR vector preparations (Fig S1). It is possible that this recombinogenic event is stimulated, in part, by sequence homology between the RBE and the p5 promoter of Rep on the pXR2 production plasmid. If so, similar AAV genomic contamination would be expected in SynITR vector preparations as the RBE element is maintained.

As many putative transcription sites were eliminated, with the potential for the creation of new ones, it was hypothesized that the transduction efficiency of AAV2-SynITR would be significantly altered. However, the data from the different human cell lines tested herein demonstrated similar transduction efficiencies to wtITR vectors (Fig. 4). These results highlight that the SynITR vectors are competent for all aspects of transduction influenced by the wtITRs, including second strand synthesis and episomal persistence, at least for the duration of those studies. Preliminary studies relying on continued cellular division in vitro and in vivo and persistent episomal forms suggest no significant difference in the ability of SynITR vectors to mediate long-term expression compared to the wtITRs. Of particular interest, the tendency of SynITR vectors to influence inadvertent host chromosome integration, is also under investigation.

The observations of similar transduction efficiencies for wtITR and SynITR vectors in cancer or immortalized cell lines, was starkly contrasted by the results following rAAV exposure to hESCs. As reported, wtITR vector transduction resulted in GFP fluorescence followed by rapid hESC apoptosis, a lethal DDR signaling event previously attributed to the wtITR sequence (2, 3). Remarkably, SynITR vectors were attenuated for induced hESC toxicity even though intracellular levels were equivalent to wtITR vectors (Fig. 5). This toxic nature of the different ITRs was emphasized by dramatic effects following AAV incubation with hESCs then induced differentiation to embryoid body formation (Fig. 6). A simple explanation for this observation is that the SynITR vectors did not uncoat in hESCs, and therefore did not induce ITR-mediated toxicity which is supported by decreased GFP cDNA and fluorescence (Fig. 5). However, a reported AAV “uncoating assay” demonstrated no significant differences between uncoated double stranded circular transgenic DNA following wtITR and SynITR vector infection (Fig. S2, (27). This result also implies that not only did the SynITRs capsids uncoat, but the transgenic genome also underwent second strand synthesis and circularized, an event known to be stimulated by the wtITR’s ability to invoke both non-homologous end joining and homologous repair pathways (32-34). Increased abundance of γH2AX, a marker of ATM activation in response to DNA double strand breaks, following treatment with both vector formats further suggests that AAV-SynITR vectors uncoated and induced upstream DDR signaling culminating in barely detectable apoptosis as noted by PARP-1 cleavage (Fig. 5C).

As previous reports demonstrated that AAV vector induced hESC toxicity was dependent upon p53 signaling, total and activated p53 protein levels were investigated (2, 3). Etoposide treatment of hESCs resulted in increased full length p53 (p53α) abundance which was also noted for cells treated with both single stranded and self-complementary wtITR vectors (Fig. 7A “N-term”). In contrast, p53α abundance did not increase following exposure to SynITR vectors (compared to the DMSO control, Fig. 7). As p53α levels increase following genotoxic stress, these results imply that SynITR vectors induce a milder DDR than wtITR vectors and presumably less toxicity (Figs. 5, 7). These p53α abundance results are supported by transactivation of the p21 promoter following wtITR vector transduction with minimal elevation exhibited in the SynITR group (113-fold vs. 4-fold, respectively (Fig. 7B)). When a marker of p53 activation was directly investigated (phosphorylation of ser15), several novel observations were observed specific to the hESC cells. First, a strong band corresponding in size to activated p53 isoforms γ or β (38kDa or 36kDa respectively) were noted specifically in unchallenged hESCs and regardless of treatment (Fig. 7A center panel). RT-qPCR using primers to detect p53γ or p53β cDNA strongly implies that the observed Western band is in fact p53β as this isoform is uniquely and actively expressed in resting hESCs over 10-fold higher than p53γ. To our knowledge this is the first report of a smaller constitutively activated p53 isoform in unchallenged hESCs, which may play a role in characterization of these cells as “primed for apoptosis” (19). When hESCs were challenged by etoposide, p53α activation was obvious as widely reported (Fig. 7A). Upon longer exposures (Fig. S4) p53α was also noted for scAAV; however, it was not seen in ssAAV wtITR or SynITR treated hESCs (Fig. S4). Interestingly, hESCs exposed to SynITR vectors demonstrated increased activation of larger p53 specific species (Figs. 7A “II”, S4) that are designated as a non-dissociated p53 homodimer in several studies (35-37). Interestingly, the p53 homodimer is thought to primarily exist in the cytoplasm and be incapable of promoter transactivation unlike the higher order nuclear tetrameric form (reviewed in (26)). This observation is under active investigation and may provide a novel hESC specific mechanism relying on the activation of differential p53 isoforms and/or varied p53 dimerization in response to rationally designed ITRs following vector transduction, and genotoxic stress in general. Finally, we want to highlight an AAV vector induced band corresponding to a small predicted p53 isoform reported in only 2 studies to date, p53ψ (25, 28). The reported function of p53ψ is to induce the expression of markers of the epithelial-mesenchymal cellular transition via a mechanism involving mitochondrial inner pore permeability (25). Therefore, p53ψ is designated as a pro-metastatic protein (25). Remarkably, p53ψ abundance increased only in hESCs exposed to rAAV, independent of the ITR, and not in hydroxyurea or etoposide treated cells (Figs. 7, S4). This novel finding requires further attention and may shed evolutionary insight into the relationship of AAV and the host DDR and provide increased clarity as to the apoptotic response to vectors containing wtITRs, yet not the SynITR vector platform.

The collective work described herein highlights the importance of the ITR sequence in transduction of particular cell types by identifying a post-entry, perhaps post-uncoating, host restriction only observed in hESCs to date. Whether similar observations will be observed in other stem cells that also undergo apoptosis upon rAAV transduction, such as neural progenitor cells and perhaps CD34+ cells, remains to be investigated (4). Furthermore, unlike most dependoviruses that contain multiple putative p53 binding sites in their ITR sequence (ITR2 contains the most at 6 sites), several pathogenic/autonomous parvoviruses do not, like the SynITR. It is tempting to speculate that if the indirect correlation between p53 binding sites in ITRs confers autonomy, then perhaps SynITR flanked wt AAV genome would demonstrate this characteristic. This hypothesis remains to be directly tested.

## Methods and Materials

### DNA Sequence Analysis

The identification of putative transcription factor binding sites in the ITR sequences relied on PROMO (Alggen) software analyses as instructed by the manufacturer. The DNA folding energy was calculated using the UNAFold Web Server based on work from Dr. Zuker (38). Putative Quadruplex forming G-Rich Sequences were predicted using the QGRS Mapper developed by the Bagga lab (39).

### Cells

Human Embryonic Kidney (HEK) 293, Huh 7, and immortalized normal human diploid fibroblast (NHF1-hTERT) cells were maintained in Dulbecco’s modified Eagle’s medium (DMEM) supplemented with 10% bovine calf serum (BCS) (Gibco) containing 100 U of penicillin and 100 mg of streptomycin (Gibco) per ml. The hESCs used herein are WiCell H9 cells (WAe009-A) and were obtained from the Human Pluripotent Cell Core at UNC (https://www.med.unc.edu/humancellcore/services-and-rates/). The H9 cells were propagated under feeder-independent conditions and maintained in an undifferentiated state using the StemFlex ™ Medium (Gibco) in Matrigel treated plates. All cells were cultured/maintained at 37°C at 5% CO_2_.

### Plasmids

The plasmids utilized in this study include a pXR2 plasmid (harboring the AAV2 *Rep/Cap* genes), the adenoviral helper plasmid pXX680 (encoding adenoviral components), and DD-based vector plasmids that harbor a CMV-green fluorescent protein (GFP) cassette (22, 40). Specifically, the wild-type ITR (wtITR) is a Holiday-like structure consisted with two copies of the D element flanking the A, B, B’, C, C’ and A’ elements of AAV2 as described previously (40). The synthetic ITR (SynITR) was designed based on the wtITR in which selected guanine-cytosine rich regions were substituted with adenine-thymine nucleotides. It was synthesized using triple ligation into the HindIII and KpnI sites of a reported pUC18 harboring a CMV-gfp cassette (20). Halves of the SynITR were produced by amplification using a long oligonucleotide as a substrate (template 1: 5’ - ATTATAGGTACCAGGAACCCCTAGTGATGGAGTTGGCCACTCCCTCTCTGCGCGCTCGCTCGCTCACTGAGGC AATTT*GATAA*AAATCGTCAAATT CGCGG-3’ and template 2: 5’-CGCGGGATAAAAATCGTCAAATTATAAACAGGCTTTGCCTGTTTAGCCTCAGTGAGCGAGCGAGCGCGCAGAGAGGGAGTGGCCAACTCCATCACTAGGGGTTCCTAAGCTTA TTATA-3’) and a respective primer set (template 1 F 5’-ATTATA*GGTAC*CAGGAACCCCTAGTGATG-3’ and R 5’-CCGCG AATTT GACGATTTTT ATC-3’; template 2 F 5’-CGCGGGATAAAAATCGTCAAATTA-3’ and R 5-TATAATA*AGCTT*AGGAACCCCTAGTGATGGAG-3’). The design produced a 5’KpnI and a 3’HindIII site for pUC18-CMV-GFP cloning and a central BsaBI site (between the two SynITR halves) that facilitated the triple ligation. DNA sequencing confirmed the final product, and BsaBI (central and internal of the SynITR) digestion was performed after every DNA preparation to ensure the SynITR integrity.

### Replication assay and analysis

The AAV DNA replication assay was performed using neutral agarose gel electrophoresis followed by Southern blotting as described previously with minor modifications (22, 41, 42). In short, 48h post-transfection (see Fig. 2 for groups), HEK293 cells were collected for Hirt DNA extraction. The cell pellet from a 10-cm plate was re-suspended in 740 µl of Hirt buffer (0.01 M Tris-HCl Ph 7.5 and 0.01 M EDTA) and lysed by adding 50 µl of 10% sodium dodecyl sulfate (SDS). The cell lysate solution was then mixed with 330 µl of 5M NaCl, incubated overnight at 4°C and centrifuged at 15,000 rpm at 4°C for 1h. The supernatant was harvested and mixed with phenol/chloroform/isoamyl alcohol (25:24:1, Sigma) and centrifuged at maximum speed (∼16,000 rpm) for 5 min. The aqueous phase (top layer) was then mixed with an equal volume of chloroform (Sigma). Low-molecular weight DNA was precipitated by the addition of an equal volume of 100% isopropanol (Thermo Fisher Scientific), followed by centrifugation at 16,000 rpm for 15 min at 4°C. The resultant pellet was resuspended in 50 µl of TE buffer (10 mM Tris–HCl, 1 mM EDTA pH 8.0) supplemented with 100 µg/ml of RNase A. The Hirt DNA yields were determined by the Nanodrop Spectrophotometer, and equivalent amounts of Hirt DNA were digested with or without *Dpn*I (New England Biolabs) and analyzed on a 0.8% neutral agarose gel followed by transfer to a nitrocellulose membrane. Detection relied on a ^32^P-labeled DNA probe specific for GFP sequence synthesized with a random priming labeling kit (Takara) followed by exposure to film and development as described (43).

### ITR sequencing

All plasmids were submitted for sequencing of the ITRs using the AAV-ITR sequencing service at Genewiz and the below primer set: F 5’-ATACCGCACAGATGCGTAAGG-3’ and R 5’-AATGGAAAGTCCCTATTGGCG-3’. For sequencing the capsid packaged ITRs, vector genomes were purified from preparations following Benzonase treatment using Proteinase K and subsequent column purification using a QIAquick PCR purification kit (Qiagen). Samples were then analyzed using Sanger sequencing.

### Vector production and titration

A standard triple plasmid transfection using polyethyleneimine (PEI)-MAX (Polysciences, MW, 40,000) method was used for AAV production as described previously (22). In short, 12 µg of pXX680 plasmid, 10 g of pXR2 plasmid, and 9 µg of vector plasmid were used per 15 cm diameter dish of 80% confluent 293 cells for vector production. Cells and the growth medium were harvested around 65h following transfection. Particles were purified using CsCl gradient ultracentrifugation and dialyzed in PBS overnight. Purified virus was stored at -80°C. The vector titers were determined by probe-based qPCR analysis in a LightCycler 96 qPCR machine following Benzonase treatment to eliminate non-encapsidated DNA(44). Two different probe/primer sets were used. One set specific to the GFP region includes the Universal probe library (UPL) probe #67 (Roche), the forward primer: 5′-GAAGCGCGATCACATGGT-3’, and the reverse primer: 5′-CCATGCCGAGAGTGATCC-3’. Another probe/primer set recognizing the ampicillin resistance gene relied upon the UPL probe #58 (Roche), the forward primer: 5’-CTAGCTTCCCGGCAACAAT-3’; and the reverse primer: 5’-GCAGAAGTGGTCCTGCAACT-3’. Plasmids and a reference AAV vector were diluted serially 1:10 in a range of 10–10^8^ copies/µl and employed as the standards. The vector yield comparisons were calculated from six independent assays. For the determination of intracellular vector genomes, including those tallied in the uncoating assay, the primer/probe set amplifying the GFP sequence (mentioned above) was used.

### B1 Western blot

To detect proteins via Western blotting, 4XSDS sample buffer was added to the virus. After denatured for 15 min at 95°C, protein were separated on 4-12% NuPAGE Bis-Tris protein gels (Invitrogen) together with a standard protein ladder (Thermo Fisher Scientific). Subsequently, proteins were transferred to a nitrocellulose membrane using the iBlot 2 Dry Blotting System (Bio-Rad) with pre-assembled dry transfer stacks. The primary antibodies B1 (reacts with AAV capsid proteins VP1, VP2 and VP3) were diluted 1:20 in blocking buffer. For detection, a horseradish peroxidase (HRP)-conjugated secondary goat anti-mouse antibody (Abcam) diluted 1:10,000 in blocking buffer was used. A luminol-based enhanced chemiluminescence (ECL) detection kit (Thermo Fisher Scientific) was used to visualize bound antibodies. Images were taken using Amersham imager 600 (GE Health). Detection of p53 or p53-ser15-P relied on Cell Signaling antibodies #2527 or #9284, respectively. The B-actin antibody was from Santa Cruz Biotechnology (sc-47778) and the JBW301clone (Sigma) was used for γ-H2AX detection.

### QPCR for transgene expression

Endogenous expression of p53 isoforms and p21 were quantitatively analyzed by qRT-PCR. Total RNA was isolated using AllPrep DNA/RNA mini kit (Qiagen), followed by DNase I treatment using DNA-free™ Kit and DNase I Removal Reagents (Catalog Number AM1906, Ambion by life technology) before reverse transcription was performed. cDNA was then synthesized with the iScript cDNA synthesis kit (BioRad) in the presence and absence of reverse transcriptase. qPCR amplification was performed with an initial denaturation step at 95°C for 10 min, followed by 45 cycles of denaturation at 95°C for 10 s, and annealing/extension at 60°C for 20 s using the following primer sets. p21 sense: TGGAGACTCTCAGGGTCGAAA, and antisense: GGCGTTTGGAGTGGTAGAAATC; P53-β sense : GCGAGCACTGCCCAACA P53-β antisense: GAAAGCTGGTCTGGTCCTGAA; P53-γ sense: TGACTTT GAACT CTGTCTCCTTCCT P53-γ antisense: GTAAGTCAAGTCAAGTAGCATCTGAAGGGTG. All reverse transcription reactions were performed using a standard PCR machine from Applied Biosystem. All qPCR reactions in this study were either performed with a Roche 480 Lightcycler or the Step One Plus instrument (Applied Biosystems).

### Quantification of episomal DNA

Cells were seeded in 24-well plates and incubated with AAV vectors, followed by total DNA extraction at 24 hrs p.i. using the Qiagen DNeasy and Blood Tissue Kit. Half of the DNA was treated with T5 exonuclease (NEB) at 37°C overnight and then 80°C for 15 min, while the remaining half of DNA was mock treated. Subsequently, the samples were diluted 5-fold, and the amount of vector genomes was quantified by qPCR using UPL probe #67 as indicated above. Virus control was included to check the efficiency and efficacy of the assay, virus was directly added into cell pellet, DNA was extracted as described above. The percentage of episomal DNA was calculated from the ratio of T5 resistant GFP DNA to total GFP DNA.

### Embryoid body formation

The EB culture was performed using DMEM-F12 (Gibco) containing 20% KnockOut™ Serum Replacement (KOSR), a final concentration of 2 mM L-Glutamine, 0.1 mM non-essential amino acids, 0.1 mM ß-mercaptoethanol, and 1X Penicillin/ Streptomycin. The medium was prepared and stored at 2-8°C for up to 2 weeks. Six hours post AAV infection, the WiCell H9 cells were rinsed with D-PBS, treated with Dispase II solution (1 mg/ml) at 37°C for 3 mins, and then gently scraped off from the culture dish. The cell clumps were then resuspended in the EB culture medium described above with the addition of ROCK-inhibitor at a final concentration of 10 µM. The medium was changed daily starting from day 2. EB plates were scanned, and the number of EB spheres were counted on Day 5 culture. The EBs were dissociated into single cells using Accutase for flow cytometry analysis or directly frozen for viral genome or cDNA analysis.

### A20 Western dot blot

The intact particles were determined by A20 western blot and densitometry analysis. In short, 2-fold serial dilutions of the virus were directly loaded onto a nitrocellulose membrane in a dot blot apparatus. Membranes were blocked in 10% milk in PBS for 30 min at room temperature and incubated with the A20 primary antibody (dilution, 1:20) in 2% milk for 1 h at RT. Membranes were washed 5 times with 1× PBST and incubated with goat anti-mouse horseradish peroxidase-conjugated secondary antibody (Abcam) for 30 mins then visualized using the chemiluminescence kit (Thermo Fisher Scientific). Image J software was used for densitometry analysis.

### Alkaline gel of AAV genome

The viral genome integrity was analyzed by alkaline agarose gel electrophoresis followed by SYBR Green I (Sigma) staining. Briefly, the purified viruses were subjected to Benzonase digestion at 37°C for 30 mins followed by heating at 95 °C for 10 mins. Prior to the loading, the viral DNA samples were mixed with EDTA at a final concentration at 50 mM and SDS at a final concentration of 1% and 1X alkaline gel-loading buffer (a 6X alkaline loading buffer contains 0.18g/ml of Ficoll, 40 mM of NaOH, 5 mM of EDTA and Xylene Cyanol)1X denaturing buffer, 10% SDS). The viral DNA samples were run in a 1 % alkaline agarose gel made with double-distilled water (ddH2O) containing a final concentration of 50 mM of NaOH and 1 mM of EDTA. The electrophoresis was performed with freshly made alkaline electrophoresis buffer (50 mM NaOH, 1 mM EDTA) at 40 v for 4 hours. After the run the gel was washed and neutralized in 1X TAE for 30 min and subsequently stained in 1x SYBR Green I nucleic acid staining solution in 1X TAE. The gel was imaged under UV light using Amersham imager 600 (GE Health).

### AAV Transduction

Cells were seeded and infected with the AAV vectors at 10,000 vg/cell, unless otherwise noted. The GFP expression were examined using Olympus microscopy at varies time points, and flow analysis was performed at 72 hours post infection for HEK293, Huh7 and NHF1-hTERT cells, or at 24 hours post infection for WiCell H9 cells due to extensive cell death observed in certain groups. Propidium Iodide (PI) was added prior the counted using the LSR Fortessa flow cytometry system and analyzed using FlowJo software (Tree Star, Ashland, OR). For WiCell H9 cells, DNA and RNA were harvested using the AllPrep DNA/RNA mini kit (Qiagen). qPCR and RT-qPCR were utilized to quantify the viral genome and mRNA levels, respectively. In short, the RNA were treated by DNAse I using a DNA-*free*™ kit (Ambion by life technologies) followed by cDNA synthesis using the iScript cDNA Synthes Kit (BioRad). The GFP transcript counts were normalized to the GAPDH transcript counts. Below primers were used for GAPDH qPCR, the forward primer: 5’-ACCCAGAAGACTGTGGATGG-3’; and the reverse primer: 5’-TTCAGCTCAGGGATGACCTT-3’. Colony sizes were monitored overtime; plates were scanned and the covered surface areas were calculated using ImageJ.

### Statistical Analyses

Statistical analyses were performed using GraphPad Prism 8 (GraphPad Software, San Diego, CA). Statistical analysis was performed using a two-way ANOVA or T test. Significant differences were defined as a p value < 0.05.

## Acknowledgements

Support was provided by the UNC’s department of Ophthalmology and by the NIH (RO1AI072176-06A1, R.J.S/M.L.H). RJS and MLH are co-inventors of the SynITR platform which is licensed to AskBIO and RJS and MLH have received royalties. MLH is a cofounder (holds equity) of Bedrock and Astro Therapeutics for which he is also a consultant, in addition to Graybug Vision. RJS is the founder and a shareholder at Asklepios BioPharmaceuticals. He holds patents that have been licensed by UNC to Asklepios for which he receives royalties. He has consulted for Baxter Healthcare and has received payment for speaking. LS is an employee of Askleopios since April 2022. The additional authors declare no conflicts of interest.

## Author Contributions

M.L.H, L.S., N.J.B., and J.J.B. performed experiments. L.S., R.J.S. and M.L.H. wrote the paper. M.L.H designed the experiments and interpreted the data. M.L.H. and R.J.S. provided funding.

---

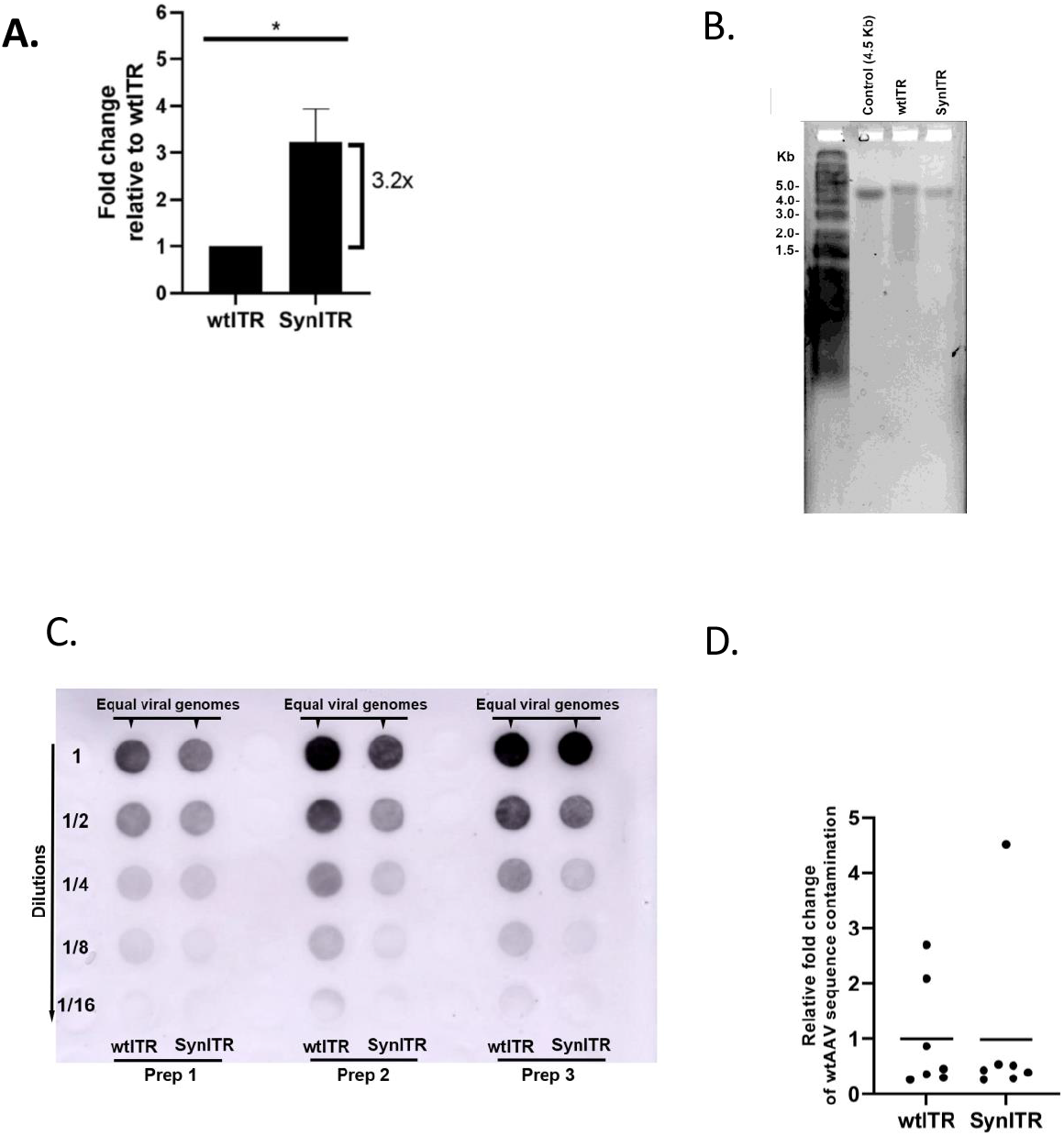

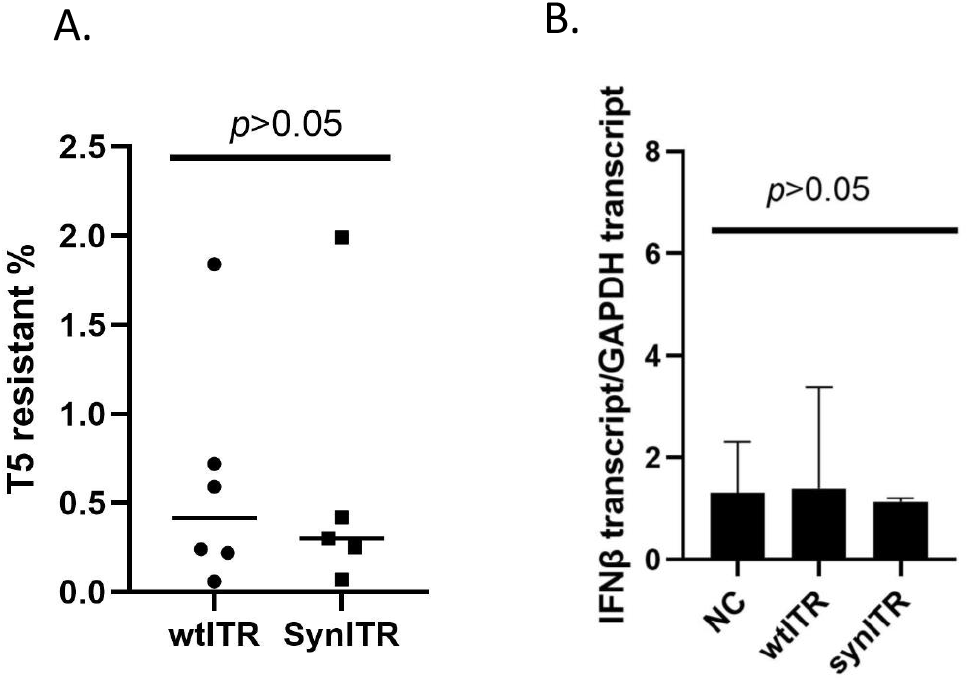

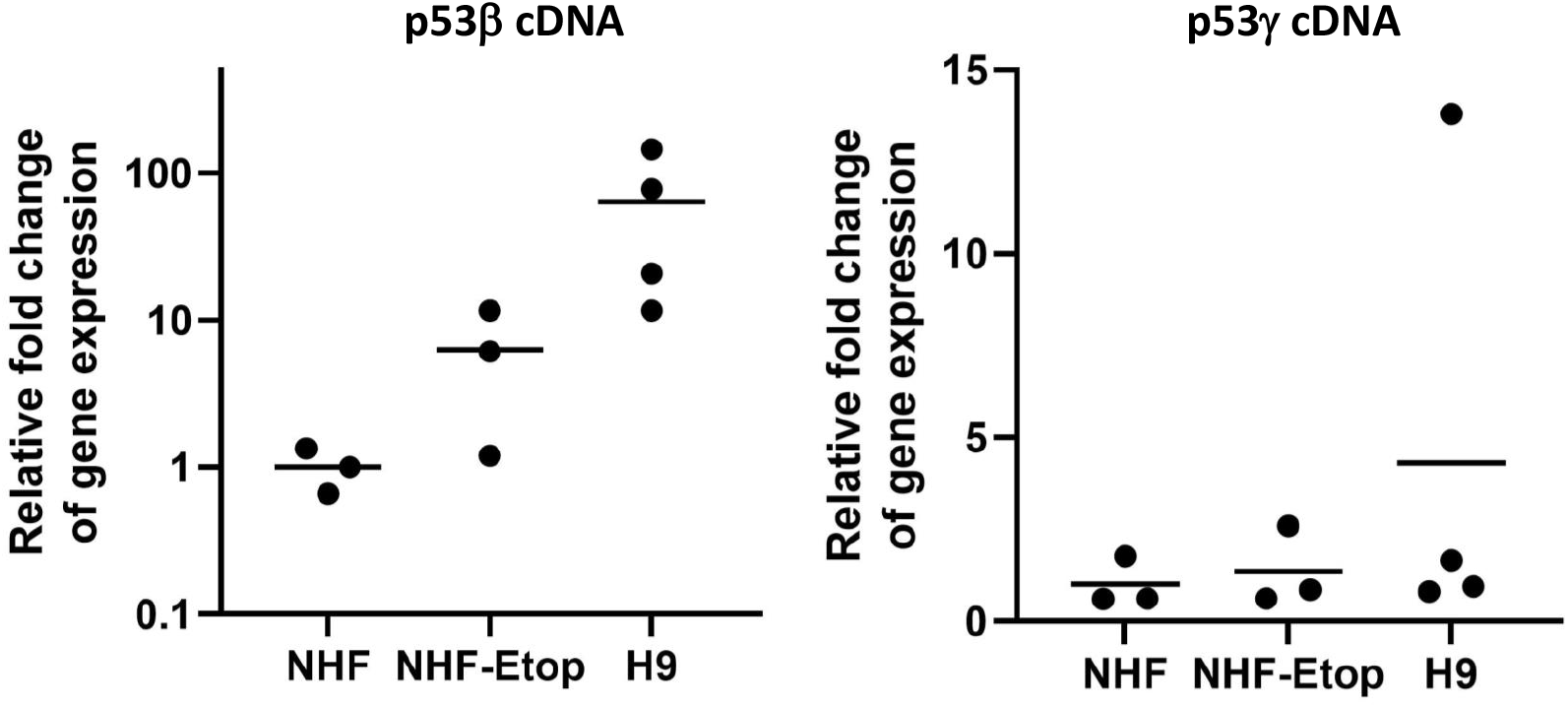

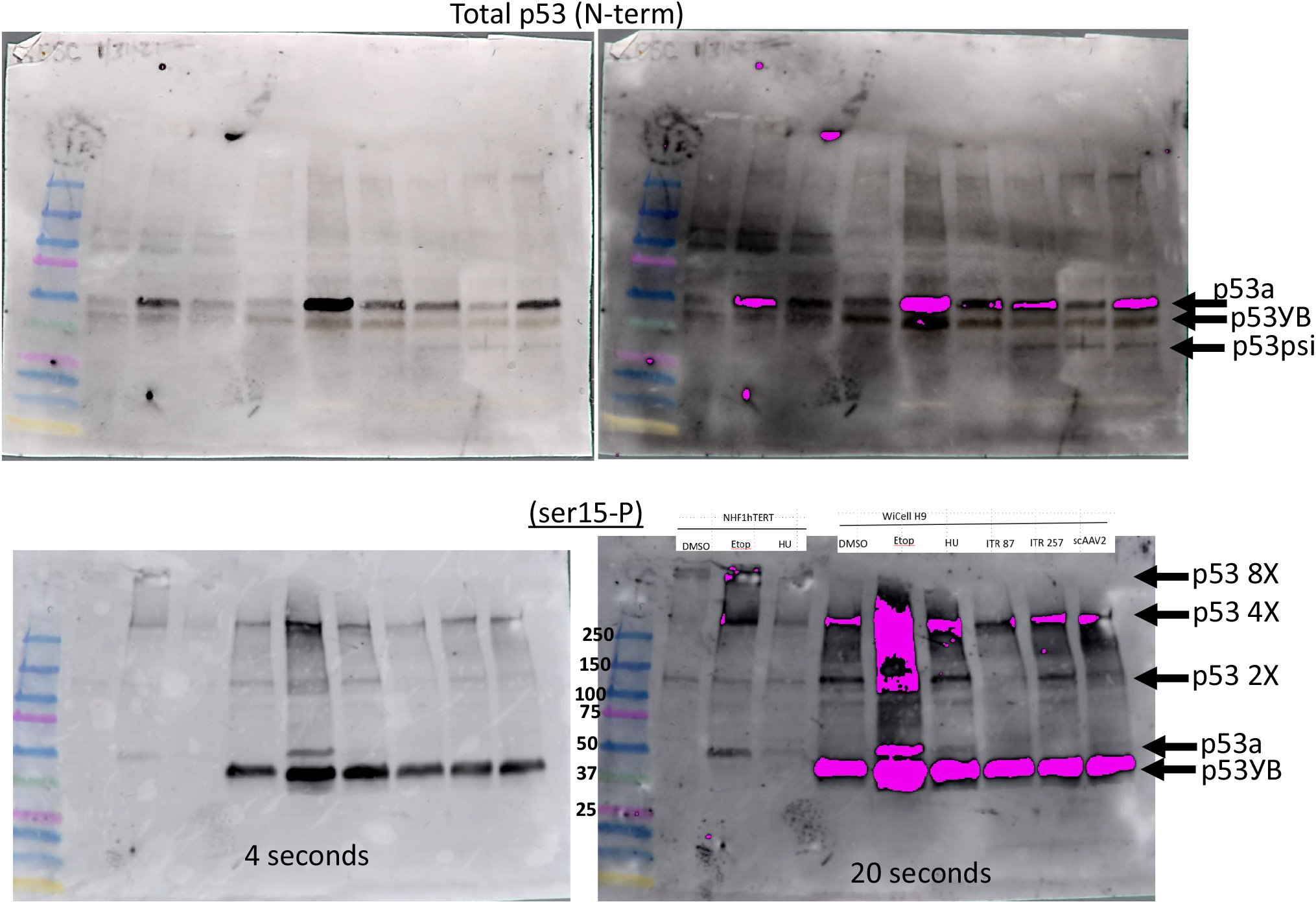

